# Likelihood-based estimation of substructure content from single-wavelength anomalous diffraction (SAD) intensity data

**DOI:** 10.1101/2021.02.07.430107

**Authors:** Kaushik S Hatti, Airlie J McCoy, Randy J Read

**Affiliations:** Cambridge Institute for Medical Research, Department of Haematology, University of Cambridge, The Keith Peters Building, Hills Road, Cambridge, CB2 0XY, United Kingdom; Drug Discovery Unit, Wellcome Centre for Anti-Infectives Research, School of Life Sciences, University of Dundee, Dow Street, Dundee, DD1 5EH, United Kingdom

**Keywords:** single-wavelength anomalous diffraction, substructure, likelihood, phasing

## Abstract

SAD phasing can be challenging when the signal-to-noise ratio is low. In such cases, having an accurate estimate of substructure content can determine whether or not the substructure of anomalous scatterer positions can successfully be determined. We propose a likelihood-based target function to accurately estimate the strength of the anomalous scattering contribution directly from measured intensities, determining a complex correlation parameter relating the Bijvoet mates as a function of resolution. This gives a novel measure of intrinsic anomalous signal. The SAD likelihood target function also accounts for correlated errors in the measurement of intensities from Bijvoet mates, which can arise from the effects of radiation damage. When the anomalous signal is assumed to come primarily from a substructure comprised of one anomalous scatterer with a known value of f” and when the protein composition of the crystal is estimated correctly, the refined complex correlation parameters can be interpreted in terms of the atomic content of the primary anomalous scatterer, before the substructure is known. The maximum likelihood estimation of substructure content was tested on a curated database of 357 SAD cases with useful anomalous signal. The prior estimates of substructure content are highly correlated to the content determined by phasing calculations, with a correlation coefficient (on a log-log basis) of 0.72.

**Synopsis:** An intensity-based likelihood method is provided to estimate scattering from an anomalous substructure considering the effect of measurement errors in Bijvoet pairs and correlations between those errors.

## 1. Introduction

The anomalous differences between Bijvoet pairs of reflections can be exploited for phasing in crystallography. However, the anomalous differences in intensities are generally limited to a few percent in size, and special care needs to be taken in planning the experiment and in collecting and processing the data in order to measure such differences with sufficient precision for successful phasing (Terwilliger *et al*., 2016*b*). Planning the experiment benefits from estimating the achievable anomalous difference, considering the number of anomalous scatterer sites that might be present and the precision to which the intensities are measured (Terwilliger *et al*., 2016*a*).

Both SHELXD (Schneider & Sheldrick, 2002) and AutoSol (Terwilliger *et al*., 2009), the experimental phasing suite in Phenix (Liebschner *et al*., 2019), require a prior estimate of how many anomalous scatterers are expected in the substructure. The most accurate estimates are obtained when there is a known stoichiometry for an intrinsically bound metal, so that the size of the substructure depends only on the number of copies in the asymmetric unit. For soaking experiments with heavy metals or halides, initial estimates of the number of sites depend on rules of thumb that are typically based on the number of residues. Even when phasing with intrinsic scatterers such as sulphur atoms in native proteins or with selenium atoms in proteins incorporating Se-Met, parts of the chain may be disordered, selenium substitution may be incomplete, or radiation damage could reduce their occupancy by the end of the X-ray diffraction data collection.

This work looks at characterising the data after the experiment has been performed and the data have been processed. Specifically, we are addressing the problem of determining, from the data, the amount of scattering contributed by the anomalous substructure. This provides both an estimate of the size of the actual anomalous differences between Bijvoet pairs and information about the number of sites that is expected in the substructure. The underlying approach is to devise a likelihood target that can be used to determine parameters that quantify the strength of anomalous scattering, considering the effect of errors in measuring intensities of Bijvoet pairs and also the effect of correlations in those errors. These parameters are also required to refine substructure models and obtain phase probability distributions from SAD intensity data. This will be investigated in future work, along with ways to assess anomalous signal through measures of information gain and estimates of the log-likelihood-gain score that would be achieved with an ideal substructure model.

### 2. Intensity-based joint probability distributions for SAD data

To derive probability distributions for measured diffraction data for use in crystallographic likelihood functions, it is necessary to combine the effects of complex differences in the structure factors with those of scalar measurement errors in the intensities. This is further complicated by the fact that the amplitude of the structure factor is related to the square root of the intensity; the true intensity is never negative, but the measured intensity may well be. We have not found a way to derive exact analytical expressions combining these differences. Nonetheless, in our previous work on the LLGI intensity-based likelihood target (Read & McCoy, 2016), we showed that a log-likelihood-gain score that accounts exactly for the effect of Gaussian measurement errors on intensities can be approximated extremely well with a target computed via the Rice function (for the acentric case), in which the intensity and its standard deviation are transformed into an effective amplitude and a Luzzati-style weighting term approximating the effect of the scalar measurement error as an error in the complex plane. Importantly, the effective amplitude and the weighting term are independent of calculated structure factors from a model, so they only need to be determined once. For the LLGI target, we assume implicitly that the intensities measured for different Miller indices are independent of each other. Here we investigate whether the same approach can be extended to intensity-based iSAD likelihood targets for SAD data, in which there is a pair of correlated intensity measurements for each set of Miller indices. We concentrate on what can be deduced about the joint distribution of the true Bijvoet mates from the corresponding intensity measurements, and what that can tell us about the scattering power of the anomalous substructure.

### 2.1. Joint prior distribution of true Bijvoet mates

To set the stage for characterising the substructure content, we start by defining the joint distribution of true Bijvoet mates (with no measurement error) in terms of the atomic content of the crystal, divided into the most significant anomalous scatterer (for which a substructure might be determined in the process of phasing) and the rest of the atoms. Note that we are not assuming here that the rest of the atoms lack any anomalous scattering contribution. For instance, in Se-Met phasing the sulphur atoms in cysteine residues will make a small but non-negligible contribution to the anomalous differences, even though it is only rarely possible to identify the positions of these atoms during substructure determination.

The Bijvoet mates are described in terms of **F**^**+**^ and the complex conjugate of **F**^**−**^, **F**^**−∗**^, because these are highly correlated and have similar phase angles. Individual elements differ in their (in general) complex scattering factor **f**, and individual atoms can differ in their occupancies and B-factors, as shown in (1a) and (1b).

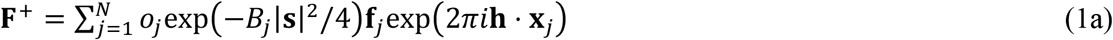

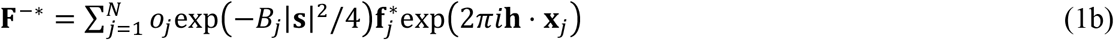

In these equations, **h** is the vector of Miller indices and **s** is the corresponding diffraction vector, the magnitude of which is the inverse of the interplanar spacing. As discussed in our earlier work on SAD phasing (McCoy *et al*., 2004), the joint distribution of Bijvoet mates takes the form of a multivariate complex normal distribution, which is readily derived by assuming that each atom contributes independently to the total structure factors. (The effects of correlations between atomic contributions arising from translational non-crystallographic symmetry (tNCS) can be addressed by modifying the expected intensity factors in the final equations, as described before for the case of normal scattering (Read *et al*., 2013)). Equation (2a) defines the prior joint distribution (before a substructure model is available), where the expected values of the complex Bijvoet mates are zero and the Hermitian covariance matrix is defined in (2b).

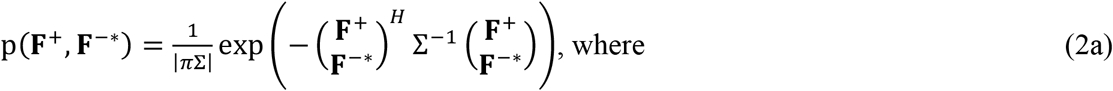

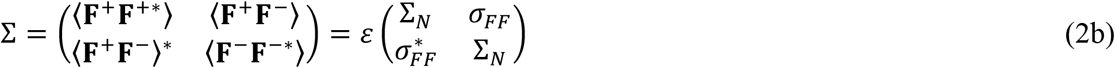

The diagonal variance term, Σ_*N*_, is simply the scattering power of the crystal defined in terms of the scattering factors in (3a), while the off-diagonal covariance element, σ_*FF*_, is defined in (3b) and ε is the expected intensity factor.

These structure factors are the sums of atomic contributions for *N* atoms, which will be divided below into the *H* atoms that could be identified as an anomalous substructure (generally a single primary anomalous scatterer type) and the remaining background (*B*) atoms that have relatively little anomalous scattering. Depending on the context, intrinsic anomalous scatterers, such as sulphur atoms in Cys and Met residues, could either comprise the *H* atoms or be considered as part of the *B* atoms if there is a stronger anomalous scatterer in the crystal. In both cases, the sums can be taken separately over the *B* and *H* subsets.

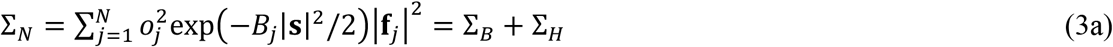

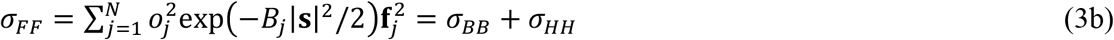

The scattering factor can be expressed as **f**_*j*_ = (*f*_0_+ *f*′) + *if*^”^, which is a function of both wavelength and resolution. Note that the wavelength-dependent correction terms, *f*^′^ and *f*^”^, are essentially independent of resolution as they arise from inner-shell electrons that can be considered to be point scatterers at relevant resolutions. The wavelength-independent form factor, *f*_0_, provides the resolution dependence. For (3a) and (3b), we can expand the scattering factor terms to obtain | **f**_*j*_ | ^2^= (*f*_0_+ *f*′)^2^ + *f*^”2^, and

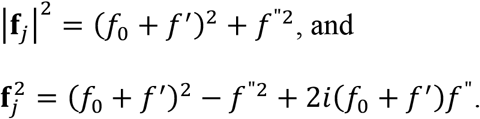

The structure factors can be normalised to give *E*-values with a mean-square value of 1. In the joint distribution of *E*-values, there is just a single complex correlation parameter, ρ_*FF*_, as shown in (4).

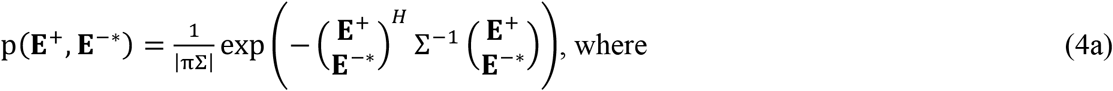

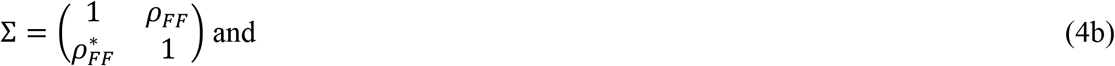

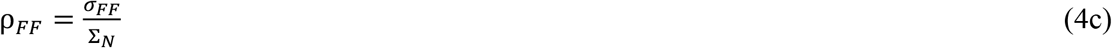

Because the Bijvoet pairs are highly correlated, values for ρ_*FF*_ in practice are only slightly less than one. The deviation from one tends to increase with resolution, because *f*^”^ is effectively independent of resolution, whereas the real parts of the scattering factors decrease with resolution.

### 2.2. Correlated measurement errors in measured Bijvoet mates

In the LLGI approach to accounting for the effect of measurement error, the intensity and its standard deviation are transformed into an effective amplitude, *F*_*e*_ (or *E*_*e*_ for normalised data) and a Luzzati-style weighting factor *D*_0_ that, in a Rice probability function, give an excellent approximation to the posterior probability of the true amplitude given the intensity. In the related iSAD approach to an intensity-based likelihood function, proposed here, both members of the Bijvoet pair are transformed in the same way.

As demonstrated below, this approach is well-justified when the measurement errors in the Bijvoet mates are uncorrelated, but requires some elaboration when they are correlated. As discussed by Garcia-Bonete and Katona (2019), time-dependent effects on the measured intensities, such as radiation damage, can lead to correlations between the errors of mean intensity measurements, and there is evidence of such correlations in some of the data sets we have examined (discussed below). Correlations in measurement errors can be accounted for by assuming that the errors are drawn from a bivariate normal distribution in which the individual variances are obtained from the data processing analysis but in which a non-zero correlation is present. For simplicity of notation, we use *Z* to represent the square of an *E*-value (or, equivalently, a normalised intensity). A joint probability distribution for the effect of correlated measurement errors on the observed normalised intensities is given in (5).

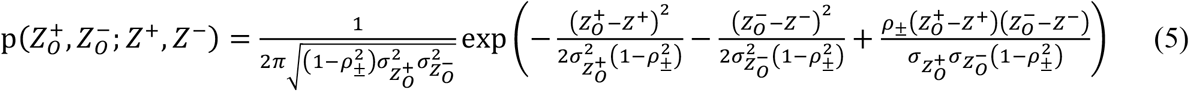

where *Z*^+^ and *Z*^−^ are the true values of the normalised intensities, 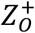 and 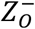 are their respective observed values, 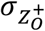 and 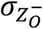 are the respective standard deviations of the measurements, and *ρ*_*±*_ is the correlation coefficient between the measurement errors.

It seems reasonable to conjecture that the effect of this correlation on the iSAD approximation can be modelled by assuming that implied complex errors in the structure factors are correlated to the same degree as the real errors in the corresponding measured intensities. In the iSAD approximation (as in the LLGI approximation), the effective normalised amplitude arises from a complex structure factor that is obtained by adding a complex normal error to the down-weighted true structure factor, as given in (6).

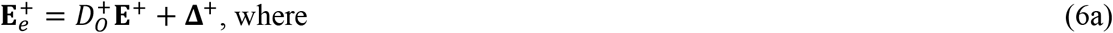

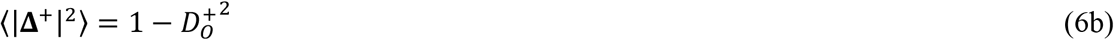

Note that, because of the 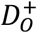 weight on **E**^+^, the expected value of 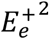 is one. Equivalent expressions apply to the Bijvoet mate. The assumption that the complex errors are correlated to each other, with a complex correlation coefficient that has a magnitude equal to *ρ*_*±*_, allows us to determine the complex correlation between 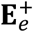 and 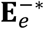, defined as *ρ*_*FF,obs*_, by analogy to the complex correlation *ρ*_*FF*_ between the corresponding true values, **E**^+^ and **E**^.−^ This is shown in (7), where we assume that the complex errors are uncorrelated with the true weighted structure factors, so that cross-terms such as 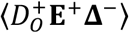 disappear.

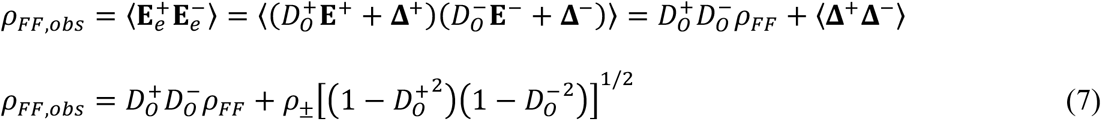

For simplicity (also justified by the consideration that the implied complex error is effectively modelling the error in the amplitude, *i*.*e*. the error parallel to the structure factor), we will assume that *ρ*_*±*_ (and thus *ρ*_*FF,obs*_) has the same phase as *ρ*_*FF*_. In any event, in the situations considered here only the absolute value of *ρ*_*FF,obs*_ influences the outcome, though the phase of the complex correlation will influence phasing calculations when a substructure model is considered in future work.

By analogy to (4), the joint distribution of the phased effective amplitudes is defined in (8).

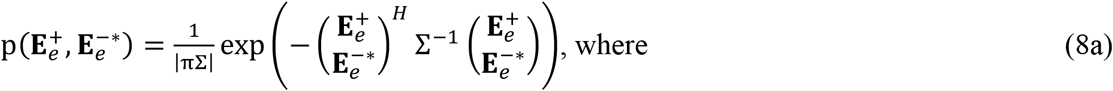

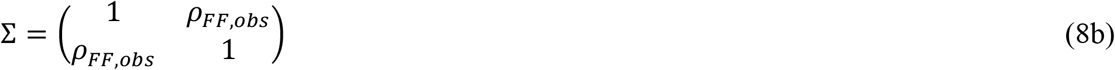

## 3. The data likelihood target: joint distribution of effective amplitudes

The probability distribution in (8) relates structure factors, but the measured data are intensities with unknown phases, which have been transformed into the effective amplitudes in this equation. The phases in (8) can be integrated out to obtain a likelihood function that depends only on the effective amplitudes, given in (9).

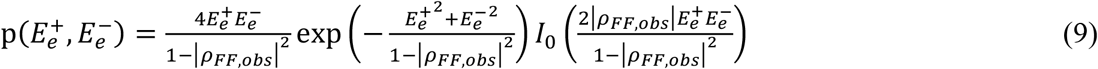

Note that there is only a single parameter to describe the variance of this distribution, *ρ*_*FF,obs*_. However, *ρ*_*FF,obs*_ is itself a function of 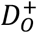 and 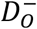, which are fixed values obtained in the calculation of the effective amplitudes, and of the adjustable parameters *ρ*_*FF*_ and *ρ*_*±*_. As discussed above, *ρ*_*FF,obs*_ can be treated as a scalar (as well as the underlying *ρ*_*FF*_) in this context, because any phase component has no effect on the likelihood in the absence of a substructure model.

This likelihood function, which is the main focus of this work, can be used for two purposes. First, the adjustable variance parameters can be refined to characterise the data in terms of the strength of anomalous scattering (|*ρ*_*FF*_|) and potentially the degree to which the measurement errors are correlated (*ρ*_*±*_), if this parameter is not available from an analysis during the merging step of data processing. Second, it provides the likelihood score for a null hypothesis in phasing, *i*.*e*. the baseline for a log-likelihood-gain (LLG) when a substructure model is available. In other words, it can play an equivalent role to the Wilson (1949) distribution in the LLG used for purely real scattering in molecular replacement or refinement. Here we will explore the uses of this likelihood function to characterise the data, particularly to estimate the substructure content.

### 3.1. Validation of iSAD approximation

To verify that it is appropriate, first, to construct the iSAD approximation by transforming the observed intensities independently into effective amplitudes and *D*_5_ factors, and second, to assume that the same correlation parameter *ρ*_*±*_ can be used to model the effect of correlated measurement error, we have followed the approach used in validating the LLGI target (Read & McCoy, 2016) by comparing the conditional probabilities of the true amplitudes given the observations, obtained either with the exact treatment or with the iSAD approximation.

The gold standard for the comparison is the joint conditional probability distribution for the true amplitudes given the observed normalised intensities, denoted as *Z*-values 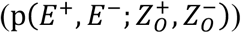 derived by following the propagation of errors and using numerical integration, giving (19) in Appendix A. The corresponding joint conditional distribution, given the effective amplitudes from the iSAD approximation 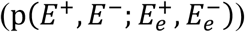, is given as (23) in Appendix B.

Fig. 1 provides two comparisons of these joint probability distributions, in a calculation modelled on Se-Met phasing where both the intrinsic anomalous signal and the measurement error are significant. In one case, the measurement errors are assumed to be independent, whereas in the second case the errors are assumed to be highly correlated with *ρ*_*±*_ = 0.75. The exact distribution and the iSAD approximation are indeed very similar for both cases, while the introduction of correlated errors has a profound effect on the distributions. Similar results are obtained in other calculations where the level of anomalous signal, measurement error and correlation of measurement error have been varied (not shown), justifying the use of this approach.

**Figure 1.**
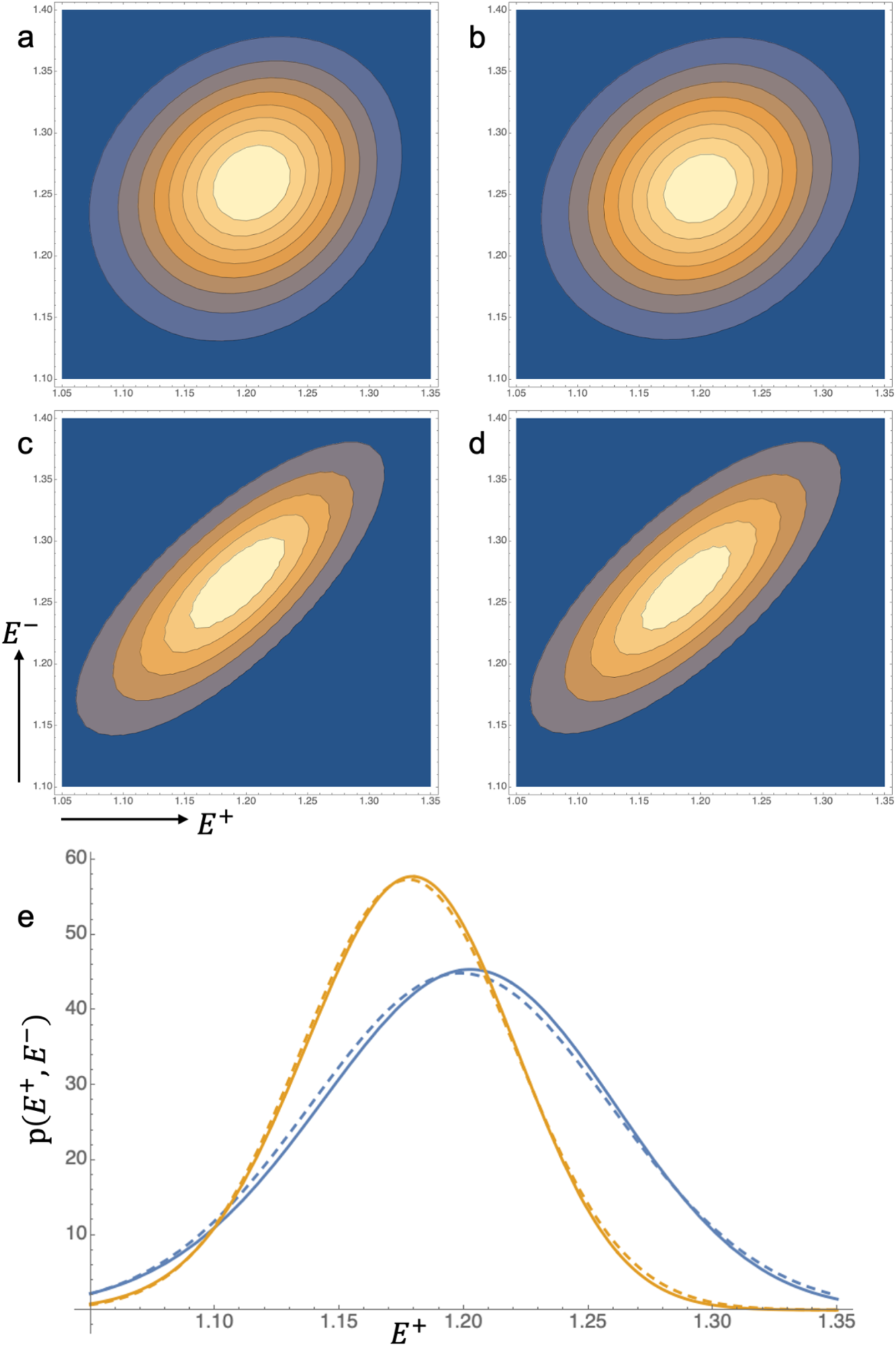
Comparison of exact and approximate probability distributions for the true normalised amplitudes conditional on the observed intensities. (*a*) Contour plot illustrating 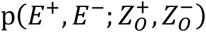 for *ρ*_*±*_ = 0. (*b*) Contour plot illustrating 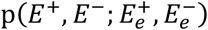 for *ρ*_*±*_ = 0. (*c*) Contour plot illustrating 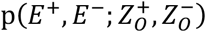 for *ρ*_*±*_ = 0.75. (*d*) Contour plot illustrating 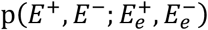 for *ρ*_*±*_ = 0.75. (*e*) Slices through the joint probability distributions at *E*^−^ = 1.25 for the cases shown in parts *a* (solid blue), *b* (dashed blue), *c* (solid orange) and *d* (dashed orange).

## 4. Maximum likelihood estimation of substructure content

When phasing with intrinsic anomalous scatterers, such as Se atoms in Se-Met constructs or S atoms in native proteins, one has reasonable prior knowledge of the atomic composition of the crystal. Even in this favourable case, there is uncertainty about the degree to which the potential sites are ordered and potential uncertainty about the occupancy of Se sites because of variable Se-Met incorporation. When soaking with heavy-atom compounds, halides or other derivatives, only a rough guess can be made in advance about the degree of substitution. Refinement of the variance parameters in a log-likelihood function based on (9) should enable a reduction of the uncertainty in the substructure content relative to other atoms in the crystal. This will be useful in characterising the phasing signal as well as in judging the difficulty of substructure determination.

There is a direct relationship between |*ρ*_*FF*_| and relative substructure content, if we treat the scattering power of only one primary type of anomalous scatterer as unknown. The anomalous scatterer content can be placed on an absolute scale if the number of copies of the protein in the crystal can be deduced from the Matthews volume (Matthews, 1968). Equation (10) is a simple consequence of (3) and (4), given that the primary anomalous scatterer (*H*) atoms share the same scattering factor, denoted **f**_*H*_ here.

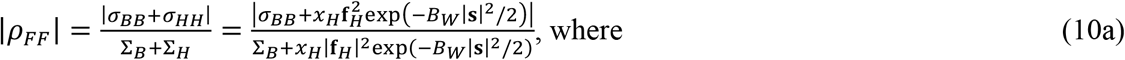

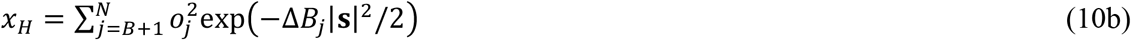

In (10a), the overall Wilson B-factor (*B*_*W*_) has been factored out of the primary anomalous scatterer contributions, leaving the individual atomic differences (Δ*B*_*j*_) in (10b). For the substructure content analysis, these equations are simplified by factoring out the overall Wilson B-factor from all sums, approximating the *B* (other background) atoms as sharing the same overall B-factor, and approximating the *H* atoms as sharing the same Δ*B*_*H*,_ relative to the B-factor of the *B* atoms. These approximations give (11).

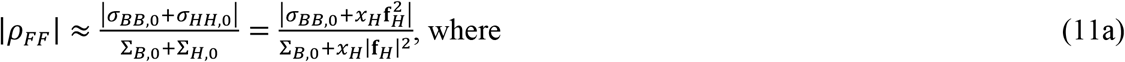

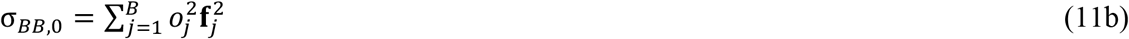

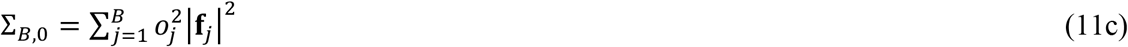

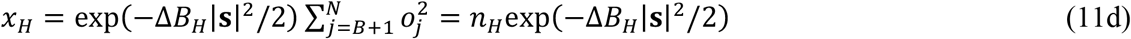

In (11d), *n*_*H*,_ is the equivalent number of fully-occupied atoms with the same total scattering power as the substructure, (which is weighted by the sum of occupancies squared).

To convert |*ρ*_*FF*_| for a resolution shell to a value of *x*_*H*_, (11a) is solved for *x*_*H*_ by transforming it into a quadratic expression in *x*_*H*_, shown in (12).

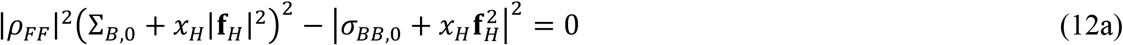

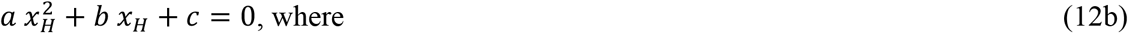

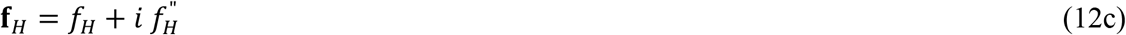

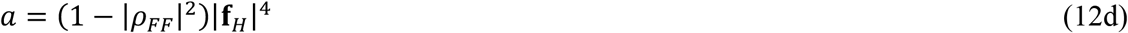

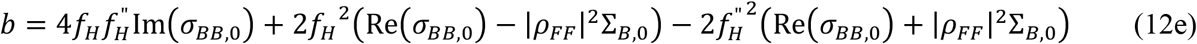

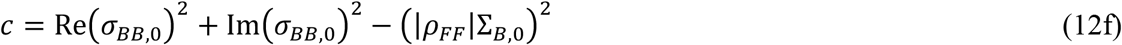

There are in general two solutions to the quadratic, as illustrated in Fig. 2. In the current implementation, the solution corresponding to a smaller substructure is chosen, though if a prior probability distribution for the substructure size were provided the two solutions could be assigned relative posterior probabilities.

**Figure 2.**
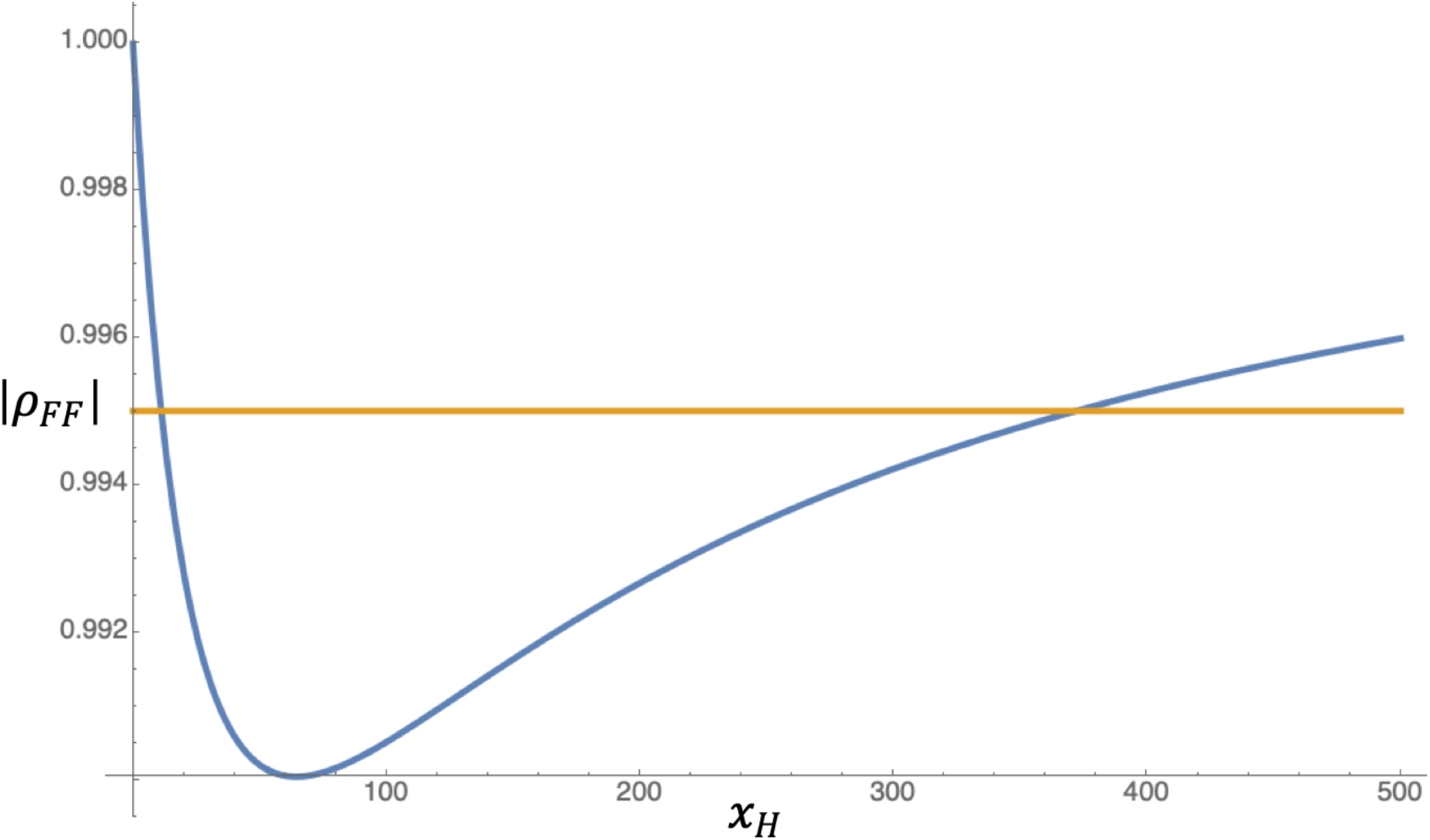
Complex correlation as a function of substructure composition. The calculated magnitude of the complex correlation, |*ρ*_*FF*_|, is shown in blue as a function of the assumed number of Se atoms in the asymmetric unit (computed against a background of 1000 C, N or O atoms and 10 S atoms). Intersections with the horizontal orange line illustrate that a refined value of 0.995 for |*ρ*_*FF*_| is consistent with either about 11 Se atoms or 370 Se atoms. The minimum value of |*ρ*_*FF*_| consistent with the assumed background composition and nature of the primary anomalous scatterer is about 0.990.

One approach that has been tested is to use the resulting *x*_*H*_ values for resolution bins to estimate values of *n*_*H*_ and Δ*B*_*H*_ by transforming (11d) into (13) and fitting a least-squares line.

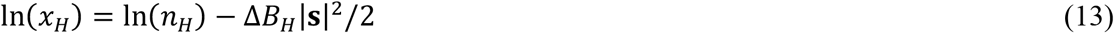

However, we have found that slightly better stability is obtained with an alternative approach. The target function given in (14) is minimized, starting from a grid search varying *n*_*H*_ and Δ*B*_*H*_ over a range of values consistent with the *x*_*H*_ estimates obtained from the refined |*ρ*_*FF*_| values.

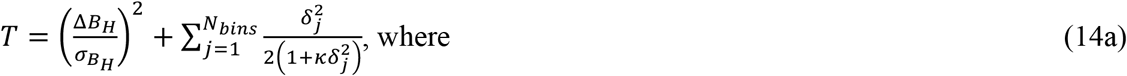

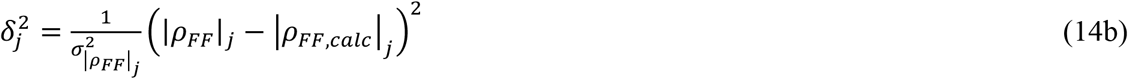

The first term in *T* restrains Δ*B*_*H*_ to zero, with a standard deviation typically set to 5Å^2^. The factor κ, typically set to 0.1, controls the damping of the robust Geman-McClure loss function comprising the second term. The calculated values of |*ρ*_*FF*_| are computed using (11). The standard deviations for |*ρ*_*FF*_| values are obtained from the inverse of the second derivative (Hessian) matrix of the likelihood target, computed for the optimised parameters.

### 4.1. Strategy for refinement of variance parameters

Refinement of the |*ρ*_*FF*_| and *ρ*_*±*_ parameters to maximise the likelihood function based on (9) is implemented in the SCA (substructure content analysis) mode of *Phasertng*, which is under development (McCoy *et al*., 2021). In the current implementation, these parameters are refined in resolution bins with a minimum of 500 reflections per bin. Two refinement macrocycles are carried out. In both macrocycles the bin values for *ρ*_*±*_ are constrained to lie in the range 0-0.9, with a weak quadratic restraint towards the value of 0 (standard deviation of 0.5) so that error correlations are inferred only when required to explain the data. In addition, a quadratic smoothness restraint penalises *ρ*_*±*_ values that differ from the value computed from the line connecting its two nearest neighbours (standard deviation of 0.025). This is similar to an approach used to stabilise the refinement of σ_U_ values for maximum likelihood refinement, when they are evaluated using just the cross-validation data (Pannu & Read, 1996). In the first macrocycle, the bin values for |*ρ*_*FF*_| are constrained to lie in the range 0-1 while being otherwise unrestrained. At the end of this macrocycle, values of *n*_*H*,_ and Δ*B*_*H*_ are estimated from the bin values for |*ρ*_*FF*_| as discussed above. Some values of |*ρ*_*FF*_| are too low to be achieved with any value of *x*_*H*_ for a given anomalous scatterer, as shown in Fig. 2. Resolution bins violating this constraint are ignored in the determination of *n*_*H*_ and Δ*B*_*H*_, and their values for |*ρ*_*FF*_| are reset to those computed from the values of *n*_*H*_ and Δ*B*_*H*_ estimated from all the data. This situation generally arises near the resolution limit, when the anomalous signal is very small relative to the noise.

For the second macrocycle, the estimated values of *n*_*H*_ and Δ*B*_*H*_ are used to determine target values for *x*_*H*_, and thus |*ρ*_*FF*_|, for each resolution shell. Loose restraints for |*ρ*_*FF*_| are applied to smooth the curve as a function of resolution, with the standard deviation being determined by the change in |*ρ*_*FF*_| that would change *x*_*H*_ by a factor of 1.5. This can stabilise refinement in cases with weak signal-to-noise but has relatively little effect in most cases. In addition, |*ρ*_*FF*_| in each bin is constrained in this macrocycle to lie between the minimum that can be achieved with any value for *x*_*H*_ and the maximum possible value, corresponding to *x*_*H*_ = 0.

## 5. Methods

### 5.1. Collecting and curating test data

The method to determine substructure content was tested on a database of SAD data sets provided by collaborators or downloaded from the worldwide Protein Data Bank (wwPDB) (Berman *et al*., 2000). 124 data sets were kindly provided by Zbigniew Dauter, most of which have been discussed earlier (Banumathi *et al*., 2004; Dauter *et al*., 2002; Wang *et al*., 2006). 162 data sets (which includes MAD data sets split into individual wavelengths and considered as SAD data sets) were collated by Tom Terwilliger from JCSG experiments and discussed earlier (Bunkóczi *et al*., 2015).

The majority of the data sets in the database were downloaded directly from the wwPDB. The advanced search option of the RCSB PDB database was used to perform queries. A list of PDB entries was collected which had a “Structure Determination Method” record containing the word “SAD”, a “Citation” record and for which experimental data including Bijvoet pairs had been deposited. Data extending to poorer than 4 Å resolution were excluded. This list was split into three categories. a) Soaking experiments comprised structures determined with any halides, heavy metals, noble gases or other elements from derivatives commonly used in phasing experiments. b) Se-Met experiments comprised structures containing selenium atoms (in order for these not to dominate the database Se-Met structures were restricted to entries deposited after 1 January 2018). c) Sulphur SAD phasing experiments were identified by examining PDB entries providing Bijvoet pairs but not containing any atoms heavier than S. For each entry, the Phenix package *phenix*.*fetch_pdb* command with the argument *--mtz* was used to download the model, sequence and structure factors, and to convert structure factors from cif to MTZ file format. The values for wavelength, cell dimensions, resolution and space group were verified with the associated publications, and any inconsistent data were removed from the list. Each data set was associated with the element type expected to contribute most strongly to the anomalous signal, denoted the primary anomalous scatterer. In total there were 536 data sets selected initially. We were surprised to note that none of these are affected by twinning or tNCS, an observation that highlights the difficulty current phasing methods have with such data.

The data sets were screened for the presence of at least minimal anomalous signal, during the initial step to generate reference substructures using the MR-SAD protocol (discussed below). Several data sets had so little anomalous signal that no anomalous scatterers could be detected. A significant number of other data sets had such poor anomalous signal that only a small fraction of atoms in the substructure were placed correctly. These data sets were omitted from subsequent analysis, leaving 382 of the original 536. It seems likely that many of these data sets are in fact native data for structures that were solved by SAD phasing using separate data that were not deposited. For a small additional number of data sets that were omitted, the reported wavelength was incompatible with the strength of the anomalous signal. This left 357 data sets in the curated database.

### 5.2. Generating reference substructures

Reference substructures were generated using the MR-SAD protocol available with *Phaser* (Read & McCoy, 2011). To be consistent in the use of structure factor amplitudes (needed for the current version of *Phaser*, which does not work with intensity data), deposited intensity data (whenever available) were converted to structure factor amplitudes (|F|) and their estimated standard deviations for the MR-SAD step, using the *phenix*.*french_wilson* tool (Liebschner *et al*., 2019). For data sets with only deposited structure factor amplitudes, these were converted to approximate intensity measurements as described for the LLGI target (Read & McCoy, 2016) for the substructure content analysis step. In the MR-SAD protocol, the deposited atomic model of the protein is used as a starting model for phasing, but treated as being composed of purely real scatterers. In the approach used here, anomalously scattering centres were found using SAD log-likelihood-gradient maps (McCoy & Read, 2010) to search for purely imaginary scatterers, since the real scattering at each centre was already accounted for in the deposited model used for phasing.

Purely imaginary scatterers found in the MR-SAD step were replaced with the atom type of the corresponding atom in the deposited structure to give the anomalous substructure, annotated using *phenix*.*emma* (Grosse-Kunstleve & Adams, 2003)(Grosse-Kunstleve & Adams, 2003) to identify atoms that superimpose within a distance threshold. Then the parameters of the anomalous substructure were refined against the data, without altering the substructure with log-likelihood-gradient completion (Read & McCoy, 2011). The refined f” for the primary anomalous scatterer and, for each anomalous scatterer type identified, the number of sites and sum of the squared occupancies of sites were stored in the database. Total scattering power of the anomalous substructure was evaluated in terms of the equivalent number of fully-occupied primary anomalous scatterers, which was calculated as the sum of squared occupancies for each atom type weighted by the square of the ratio of the f” for that anomalous scatterer type and the f” of the primary anomalous scatterer. This approximation assumes that the contribution of any secondary anomalous scatterers, if present, is dominated by their imaginary contribution, and that differences among atom types in the ratio of real to imaginary scattering are less important. Quality of SAD phasing was assessed by computing the correlation between the experimentally-phased map and density generated from the deposited model, using *phenix*.*get_cc_mtz_pdb*.

### 5.3. Choice of refined f” over theoretical f” for estimating anomalous signal

Many of the test data sets have been measured at a wavelength near the absorption edge of the primary anomalous scatterer, where the f” changes rapidly. For these data sets, the f” for the primary scatterer was refined as part of substructure refinement and phasing. Values of f” obtained from table lookup can have significant errors: tabulated values do not account for effects of the chemical environment (Evans & Pettifer, 2001), and the wavelength may not be known precisely because of errors in monochromator calibration (Ruslan Sanishvili, personal communication). It is best to obtain prior estimates of f” from a fluorescence scan of the crystal at the beamline (Evans & Pettifer, 2001), but in this study we do not have access to fluorescence scan data for the test data sets. For the data sets collected near the absorption edge, we have therefore used the refined f” values for the primary anomalous scatterer obtained during refinement and phasing with the reference substructure. We expect the refined f” to be a better estimate for the true f” than the value from a table lookup, but there will be random errors. In the refinement, the f” value for an anomalous scatterer type and the overall occupancies of the individual atoms will be correlated, with both changing the imaginary terms in calculated structure factors but differing in how they affect the relative contributions of the real and imaginary terms as a function of resolution; how well these effects are decoupled will depend on the precision of the data. There may also be systematic errors. For instance, if there is a mixed substructure and some atom types are incorrectly identified, the refined f” values will reflect a compromise between the relative real and imaginary scattering of the different atom types.

### 5.4. Preparation of data for substructure content analysis (SCA)

The diffraction data were processed using the *Phasertng*.*xtricorder* module of *Phasertng* (McCoy *et al*., 2021). *Phasertng*.*xtricorder* carries out a series of data analyses to detect and correct for the statistical effects of anisotropy, tNCS and twinning, though none of the data included in this study were affected by either tNCS or twinning. The data were scaled and used for maximum likelihood estimation of substructure content, which is carried out within *Phasertng*.*xtricorder* when the data include Bijvoet pairs. The known protein composition of the crystal was used when scaling the data; if an incorrect composition were used, the intensity scaling and therefore the estimated anomalous scatterer content would change proportionally.

### 5.5. Analysis of effect of radiation damage

To test the hypothesis that positive correlations between the measurement errors for members of the Bijvoet pair can arise from the effects of radiation damage, we searched the Integrated Resource for Reproducibility in Macromolecular Crystallography (Grabowski *et al*., 2016) (IRRMC; http://proteindiffraction.org/) to find a data set with the keyword “SAD”, strong anomalous signal, and high redundancy so that subsets of the full data could be analysed. The search yielded the data for PDB entry 3ot2 (Joint Center for Structural Genomics, unpublished) with accession identifier DOI:10.18430/M33OT2. The data set comprises 360 images, which were integrated using XDS (Kabsch, 2010) from the XDSGUI (https://strucbio.biologie.uni-konstanz.de/xdswiki/index.php/XDSGUI). Subsets of the integrated data were scaled and merged in the same package, before comparing the values obtained for the error correlation parameter as a function of resolution.

All the calculations were performed on a Dell precision 5820 machine with 128 GB RAM and an Intel Xeon W-2145 CPU @ 3.7GHz x 16, running the CentOS version 7 operating system.

## 6. Results

### 6.1. Overview of curated database

The curated database consisted of 357 data sets for crystals representing a total of 23 different anomalous scatterers (Table 1). In 22 cases, a mixture of anomalous scatterer types contributes strongly (with secondary anomalous scatterers contributing up to 50% of the total anomalous scattering). The space group sampling of the database is similar to the space group sampling of the wwPDB (Wukovitz & Yeates, 1995). Of the 357 data sets, 251 had intensity data deposited while the rest had structure factor data alone. Fig. 3 shows distributions for a number of characteristics of the data. The resolution of the data sets ranges from 0.93 Å to 3.6 Å with the total anomalous scattering ranging from the equivalent of 0.05 to 134 fully-occupied atoms. The database included data collected across a range of wavelengths from 0.81 Å to 2.29 Å; the largest peak in the distribution of wavelengths includes 143 Se-SAD data sets collected near the Se absorption edge of about 0.98 Å. There are three other notable peaks in the wavelength distribution, one near 0.9 Å largely corresponding to high-energy-remote Se data, one near 1.3 Å corresponding to the Zn absorption edge and one at 1.5418 Å corresponding to CuKα home sources. The map-to-model correlations range from values around 0.2 up to 0.8 for data sets with very high anomalous signal.

**Table 1.** Number of entries for each of the anomalous scatterers present in the database.

| Number of entries | Atom Type |
| --- | --- |
| 196 | Se |
| 58 | Zn |
| 26 | I |
| 22 | S |
| 12 | Ca |
| 9 | Hg |
| 7 | Br |
| 5 | Fe |
| 4 | Au |
| 3 | Cd |
| 2 | Ag, Mn |
| 1 | As, Ba, Co, Cu, Pt, Rb,<br>Ta, Tb, V, W, Yb |

Note: The total number of entries is greater than 357 as some data sets are counted multiple times when they contain more than one type of anomalous scatterer.

**Figure 3:**
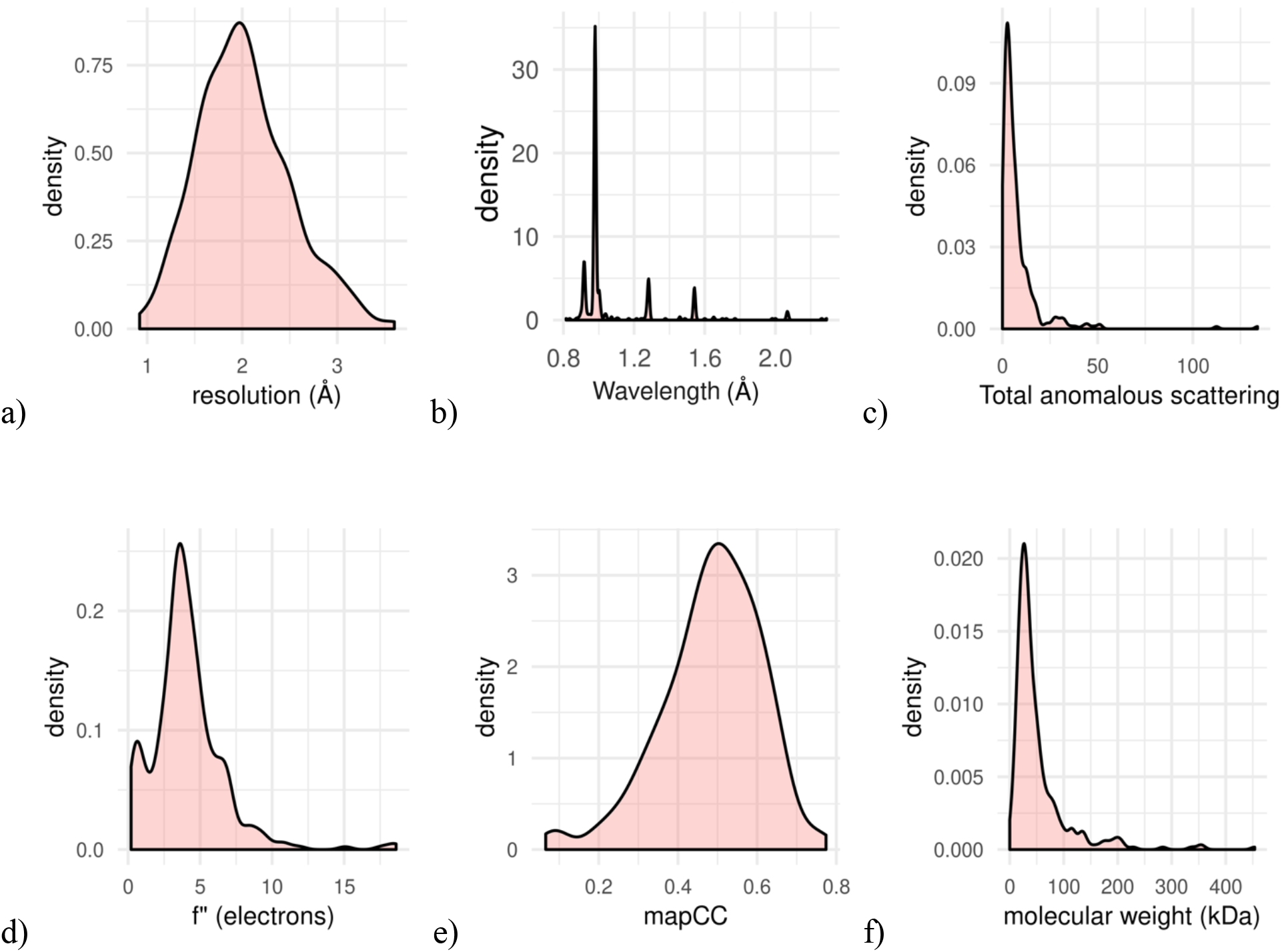
Distributions of relevant characteristics of data sets in the database. The vertical axes represent kernel density distribution. (*a*) Distribution of resolution limits; data to worse than 4 Å resolution were excluded. (*b*) Distribution of wavelengths. (*c*) Distribution of total anomalous scattering for the reference substructures, corresponding to the number of fully-occupied anomalous scatterers with equivalent scattering power. This is measured as the f”-weighted sum of squared occupancies of refined sites to account for both primary and secondary anomalous scatterers. (*d*) Distribution of refined f” values after the log-likelihood-gradient completion protocol. (*e*) Distribution of correlation coefficients between the experimentally-phased map at the end of the log-likelihood-gradient completion protocol and density corresponding to the deposited model. (*f*) Distribution of molecular weights of the target proteins.

### 6.2. Estimation of total anomalous scattering

The SCA mode estimates the scattering power of the anomalous substructure, measured in terms of the equivalent number of fully-occupied primary anomalous scatterers as discussed in section 3. The estimated number correlates well with the total anomalous scattering determined from the reference substructure, with a log-log correlation coefficient of 0.72 for data deposited as intensities (Fig. 4). (Supplementary Fig. 1 shows the equivalent plot including also data deposited as amplitudes; the correlation coefficient is still 0.72 but there are more outliers, likely reflecting the difficulty in reversing the transformation from intensities to amplitudes.) The estimates are also consistent across different element types (see supplementary Fig. 2). However, the total anomalous scattering tends to be slightly underestimated (or, alternatively, the refined occupancies could be slightly overestimated).

**Figure 4.**
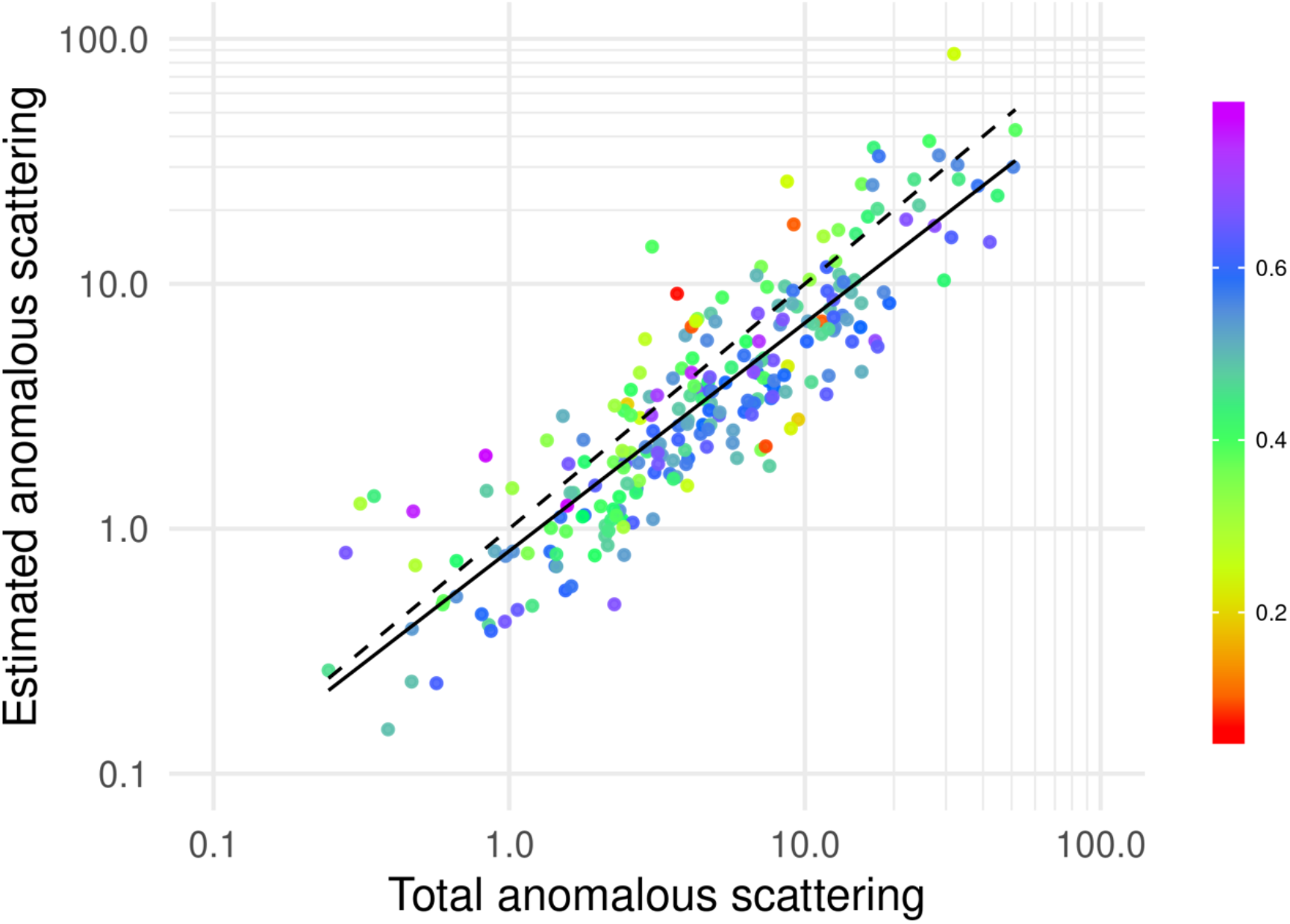
Estimation of equivalent fully-occupied number of primary anomalous scatterers, for data deposited as intensities. The horizontal axis is the total anomalous scattering power of the gold-standard substructure (weighted sum of squared occupancies of refined sites) and the vertical axis is the estimated anomalous scattering power. The dashed black line represents a perfect prediction while the black line shows the least-squares linear fit of the estimates. Each data point is coloured by the map correlation coefficient as shown in the legend. Both axes are plotted on a log_10_ scale.

### 6.3. Effects of radiation damage

PDB entry 3ot2 is in the cubic space group P23, and the diffraction data deposited in the IRRMCC comprise 360 images with 0.5° oscillation per image, giving a total of 180° of data. With the high symmetry, there is greater than 10-fold average redundancy for each observation of the plus or minus hand of the Bijvoet pairs. To confirm the presence of radiation damage during data collection, a model-phased difference Fourier was computed, comparing the data processed from the first 90 images with those from the last 90 images. The strongest peaks in the resulting map reveal decarboxylation of a number of acidic side chains, a cluster of which are shown in Fig. 5.

**Figure 5.**
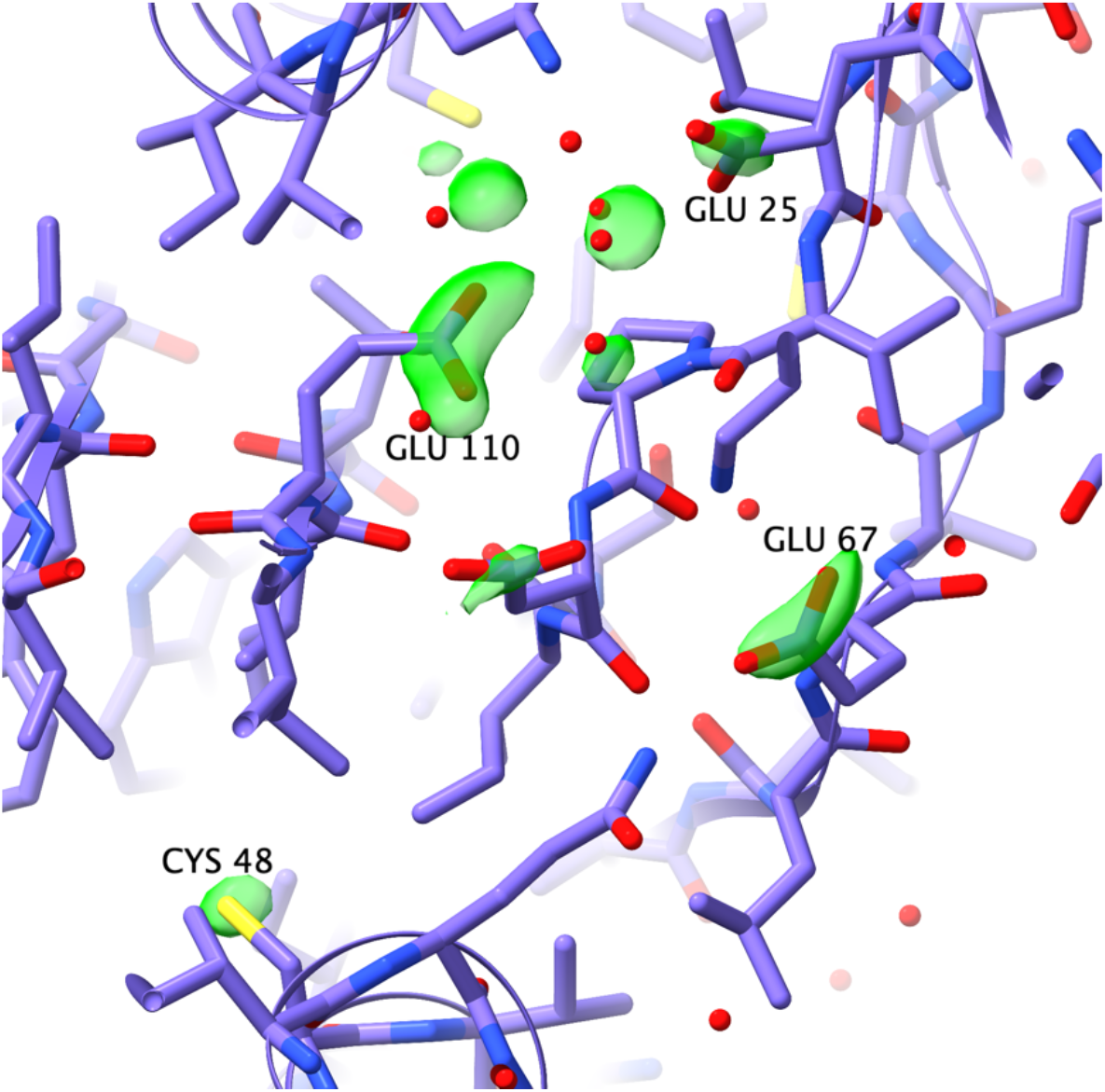
Difference map between data from the first quarter and the last quarter of data collection for PDB entry 3ot2, computed with map coefficients *F*_*O,first*_ − *F*_*O,last*_, *α*_*calc*_, with phases calculated from the deposited structure. The map is contoured at 5 times its rms value; the strongest peak, at residue Glu110 of chain A, has a height of 8.41 times the rms. This figure was made with ChimeraX (Goddard *et al*., 2018).

The diffraction data were reprocessed to include four progressively wider ranges of radiation dose, including the first 90, 180, 270 or all 360 images. The substructure content analysis was carried out for each merged data set, comparing the values of *ρ*_±_ obtained in each analysis. As expected from the hypothesis that a correlation of errors between Bijvoet mates can arise from merging data suffering from different degrees of radiation damage, the values of *ρ*_±_ increase with both resolution and total radiation dose (Fig. 6). The overall mean values of *ρ*_±_ are 0.086 for data from the first 90 images, 0.122 for the first 180, 0.149 for the first 270 and 0.160 for all 360 images.

**Figure 6.**
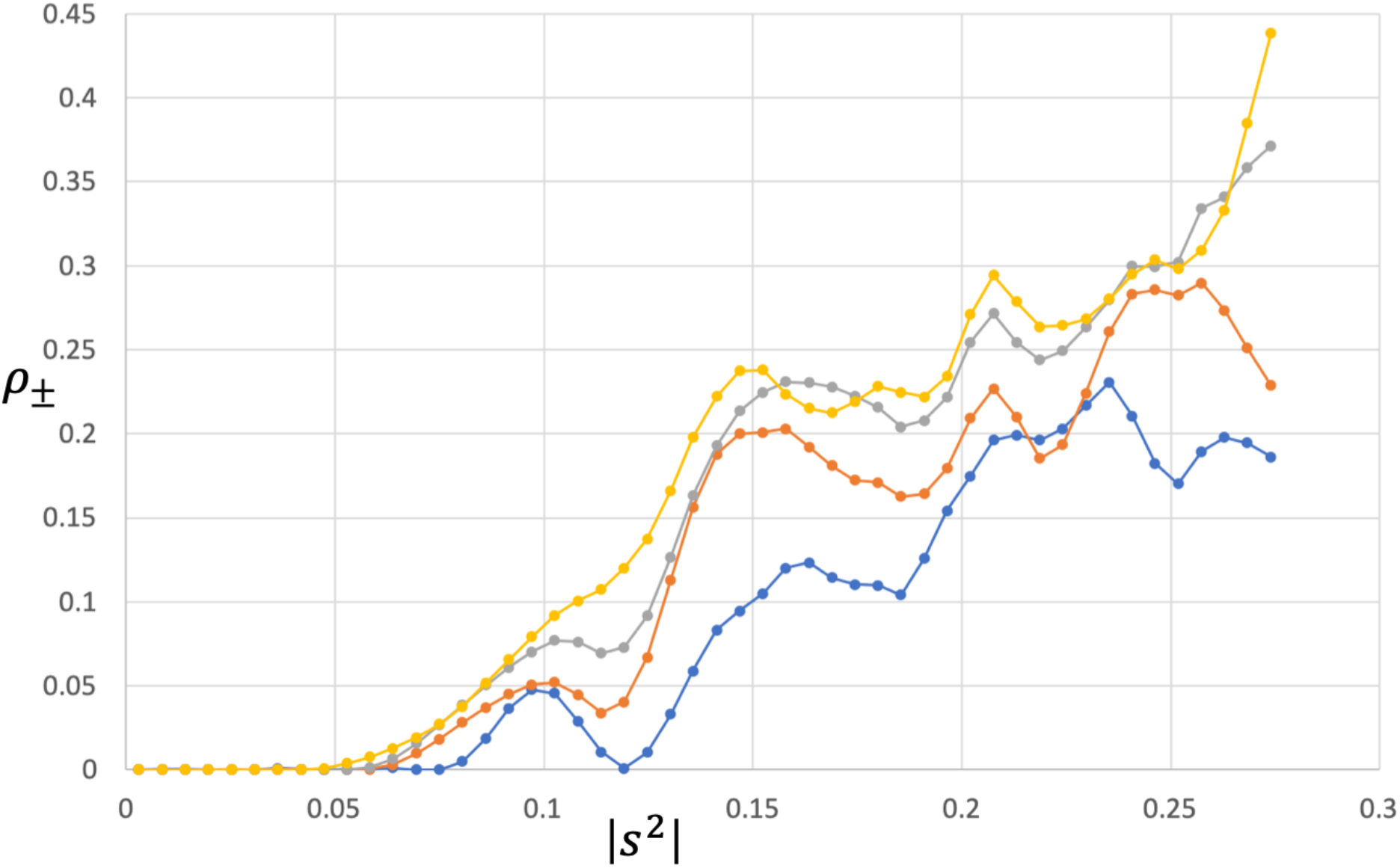
Error correlation parameter, *ρ*_±_, as function of resolution for different total levels of radiation dose. The curves show values obtained from analysing data merging the first 90 images (blue), the first 180 (orange), the first 270 (grey) and all 360 (yellow).

## 7. Discussion

In SAD phasing based on structure factor amplitudes, the difficulty of reliably extracting the anomalous signal from the noise introduced by intensity measurement errors is further complicated by difficulties in converting intensity errors into amplitude errors. Our experiences with accounting for the effect of intensity measurement errors in molecular replacement (Read & McCoy, 2016) suggested that the effects of scalar errors in intensity measurements could be approximated well as complex errors in structure factors, transforming the intensity data into effective amplitudes (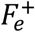 and 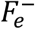) and Luzzati-type weighting parameters (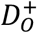 and 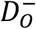). Numerical tests showed that the joint distribution of the true amplitudes in the Bijvoet pair, given the observed intensities, was approximated extremely well by this treatment, when the measurement errors in the Bijvoet pair are independent. However, the results of preliminary test calculations suggested that, in fact, measurement errors are positively correlated. A measurement error correlation parameter, *ρ*_±_, was introduced and further numerical tests showed that the joint distribution of the true amplitudes could still be approximated extremely well, even with strongly correlated measurement errors. This error treatment, therefore, will underlie our continuing work on an intensity-based SAD likelihood target, termed iSAD, which should strengthen the use of SAD data sets with marginal signal-to-noise. The joint distributions of Bijvoet mates require knowledge of the atomic composition of the crystal and the atomic scattering factors (including the anomalous, or imaginary, contributions), which is generally only known approximately when collecting diffraction data from a crystal. The role of atomic composition in anomalous scattering can be summarised by a complex correlation parameter, *ρ*_*FF*_, which varies smoothly with resolution and can therefore be determined in resolution shells. The joint distribution of observed amplitudes that takes account of the effects of anomalous scattering (*ρ*_*FF*_), measurement error (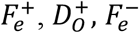 and 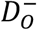) as well as the correlations in measurement errors between Bijvoet pairs (*ρ*_±_) is the basis for a likelihood target that can be optimised in terms of the two types of correlation parameters, *ρ*_*FF*_and *ρ*_±_. Given the atomic composition of the protein component of the crystal, as well as the presumed identity of the primary anomalous scatterer, the variation of *ρ*_*FF*_with resolution can be interpreted in terms of the content of the primary anomalous scatterer (equivalent number of fully-occupied atoms) and the average difference between the B-factors of the anomalous scatterers and other atoms in the crystal. In practice, if different hypotheses about the number of copies of the protein in the asymmetric unit were being tested, the estimated anomalous scatterer content would change proportionally.

The validity of the likelihood target and the deductions it allows about the anomalous scatterer content was tested by carrying out calculations on our extensive curated database. This demonstrated an excellent correlation between the predicted anomalous scatterer content and the content obtained by refining the known substructures against the same data.

The results presented here demonstrate the accuracy of the new statistical model for the effects of measurement error and atomic composition on the measurement of Bijvoet pairs of reflections. The deduced anomalous scatterer content can inform strategic decisions about whether it is likely that the substructure can be determined, how difficult the problem will be (as it depends strongly on the number of atoms to be found), and how to approach the substructure determination. Success of the statistical approach depends on the quality of the measurement error estimates; our results imply that those error estimates, at least for data used successfully for SAD phasing, are reasonably accurate.

Work in progress will build on what is presented here, showing that the results of the substructure content analysis can subsequently be used to calculate a number of measures of signal for SAD phasing: the extra information content gained by measuring Bijvoet pairs, and expected values for the log-likelihood-gain, figures of merit, and map correlations that will be achieved in phasing once a substructure has been determined. In the longer term we plan to implement a new iSAD phasing calculation, which should yield better quality phase information for data with low signal.

## Acknowledgements

This research was supported by funding from CCP4 (KSH), a Wellcome Trust Principal Research Fellowship (RJR: grant 209407/Z/17/Z) and the NIH (grant P01GM063210 to RJR), which is gratefully acknowledged. We thank Tom Terwilliger and Zbyszek Dauter for kindly sharing diffraction data sets used in this study.

## Appendix A Conditional probability of true amplitudes given the observed intensities

The first step in deriving the desired conditional probability distribution is to obtain the joint prior distribution of true normalised amplitudes, starting from the joint distribution of normalised structure factors in (4). A change of variables to amplitudes and phases yields (15), where *Z* denotes the square of the corresponding *E* value and, for notational simplicity, the phase of **E**^−*^ is referred to as *α*^−^.

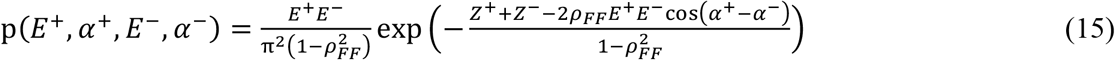

In (15), the phase of *ρ*_*FF*_has been ignored; if it were included, it would simply add a phase shift to the phase difference and would therefore have no effect on the integral over all phases in the next step to obtain the joint distribution of normalised amplitudes in (16).

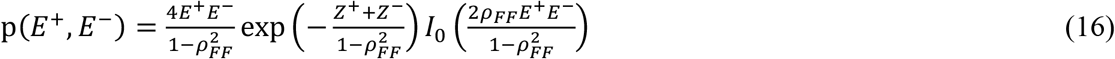

A change of variables gives the prior joint distribution of the normalised intensities in (17).

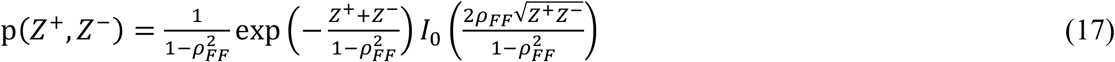

Assuming that the measured normalised intensities are related to the true values by the addition of correlated measurement errors drawn from a bivariate normal distribution, the conditional distribution of the observed normalised intensities is given in (5), repeated here for convenience.

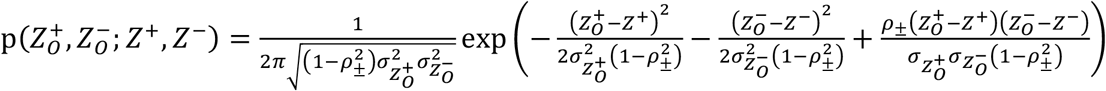

The joint probability of both pairs of true and observed intensities, (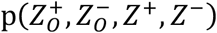*Z*^+^, *Z*^−^), is obtained by multiplying together the expressions in (17) and (5), and then the probability distribution of the observed pair of intensities is obtained by integrating over all possible values of the true intensities, shown in (18).

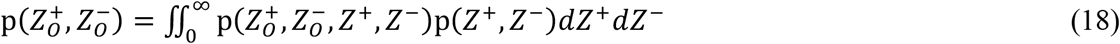

Finally, Bayes’ theorem is used to obtain the conditional probability of the true pair of intensities given the observed pair from the expressions in (5), (17) and (18), and a change of variables gives the probability distribution for normalised amplitudes shown in (19).

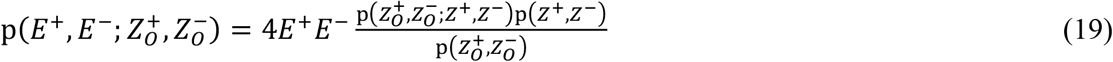

In the evaluation of (19) used for numerical tests, the double integral from (18) is carried out analytically in *Mathematica* (Wolfram Research; Inc., 2019).

## Appendix B Conditional probability of true amplitudes from the iSAD approximation

The first step in developing the desired probability distribution is to construct the joint distribution of the true normalised structure factors along with the phased structure factors corresponding to the effective amplitudes. The mathematical form for the relationships among these structure factors is the same as that considered for phasing SAD data when there are calculated structure factors from a substructure model, so the derivations below follow a similar outline to previous work on the SAD likelihood target (McCoy *et al*., 2004). To define the distribution, we need four new complex covariances (equivalent to complex correlations, because the variables are normalised), given in (20).

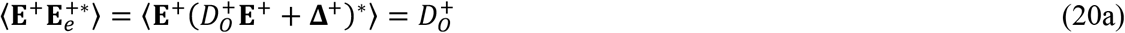

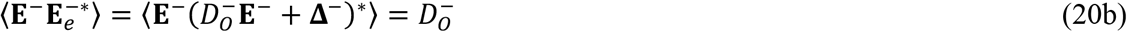

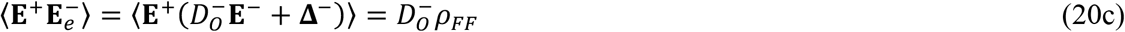

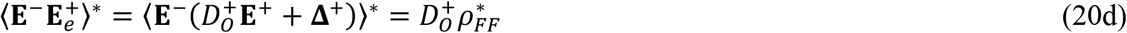

The overall joint distribution of these four complex structure factors is a multivariate complex normal distribution (21a) in which the expected values (before any measurements or other information) are all zero and the covariance matrix is given in (21b).

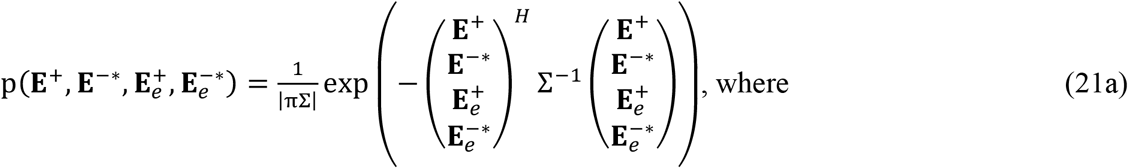

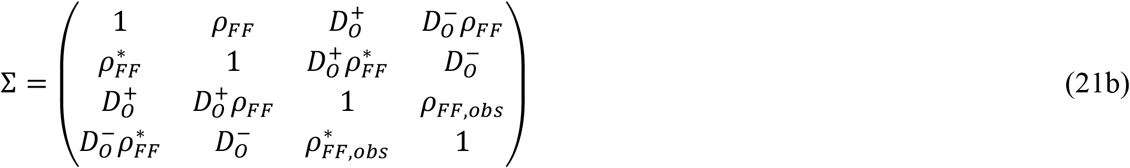

The basic strategy to obtain the desired conditional distribution is to change variables to amplitudes and phases, integrate over all the unknown phases to get a joint distribution of the amplitudes, and then apply Bayes’ theorem. One route to this result is given in (22).

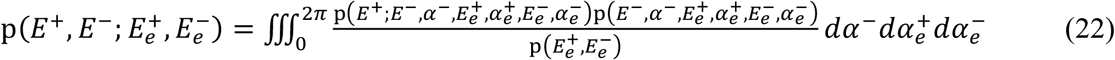

In the triple integral, all the phase terms contain phase differences so, if one phase is fixed at an arbitrary value, the others will vary over all possible values relative to each other and to the fixed phase. For this reason, one of the phase integrals can be omitted and the remaining double integral can simply be multiplied by 2*π*, as shown in (23).

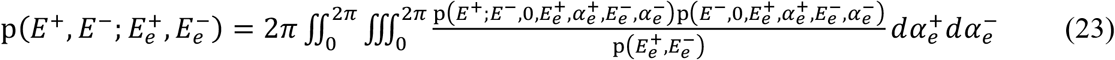

For a similar reason the phases of *ρ*_*FF*_ and _*ρFF*,obs_ are ignored, because they would just add a constant offset to the phase differences. The double integral is carried out analytically in *Mathematica* for the numerical tests. The three probability distributions needed for (23) are provided below.

The joint probability distribution for three phased structure factors, 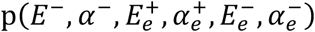), is obtained by analogy to (21), but omitting **E**^+^ as well as the first row and column of the covariance matrix and then changing the complex variables to amplitude and phase variables.

The probability of the amplitude of *E*^+^ given the other three phased structure factors is obtained by first partitioning the covariance matrix from (21) to get the conditional distribution of **E**^+^ in (24).

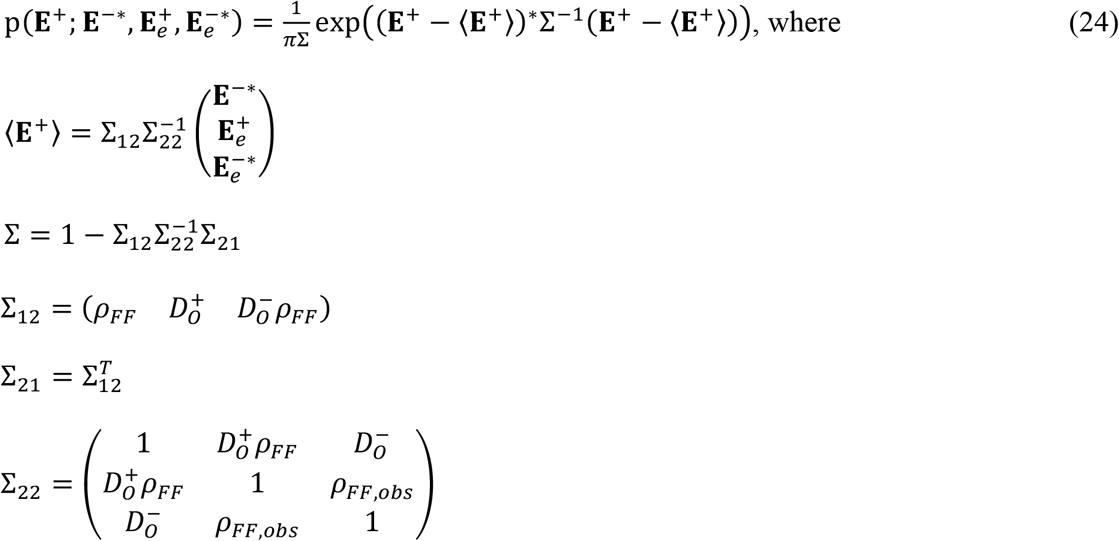

Next, after a change of variables from the complex **E**^+^ to its amplitude (*E*^+^) and phase (*α*^+^), the expression in (24) is integrated over all possible values of *α*^+^. This integral has an analytical solution, given in (25).

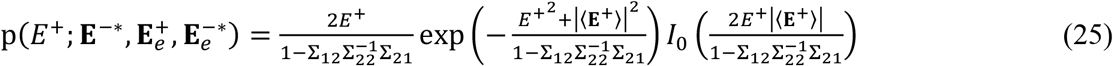

Finally, the prior joint probability distribution of the effective amplitudes, (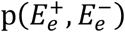), is given in (9) above, repeated here for convenience.

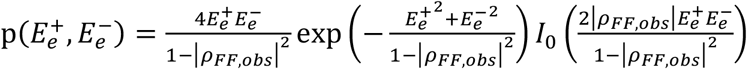

## Supplementary material

**Supplementary Figure 1.**
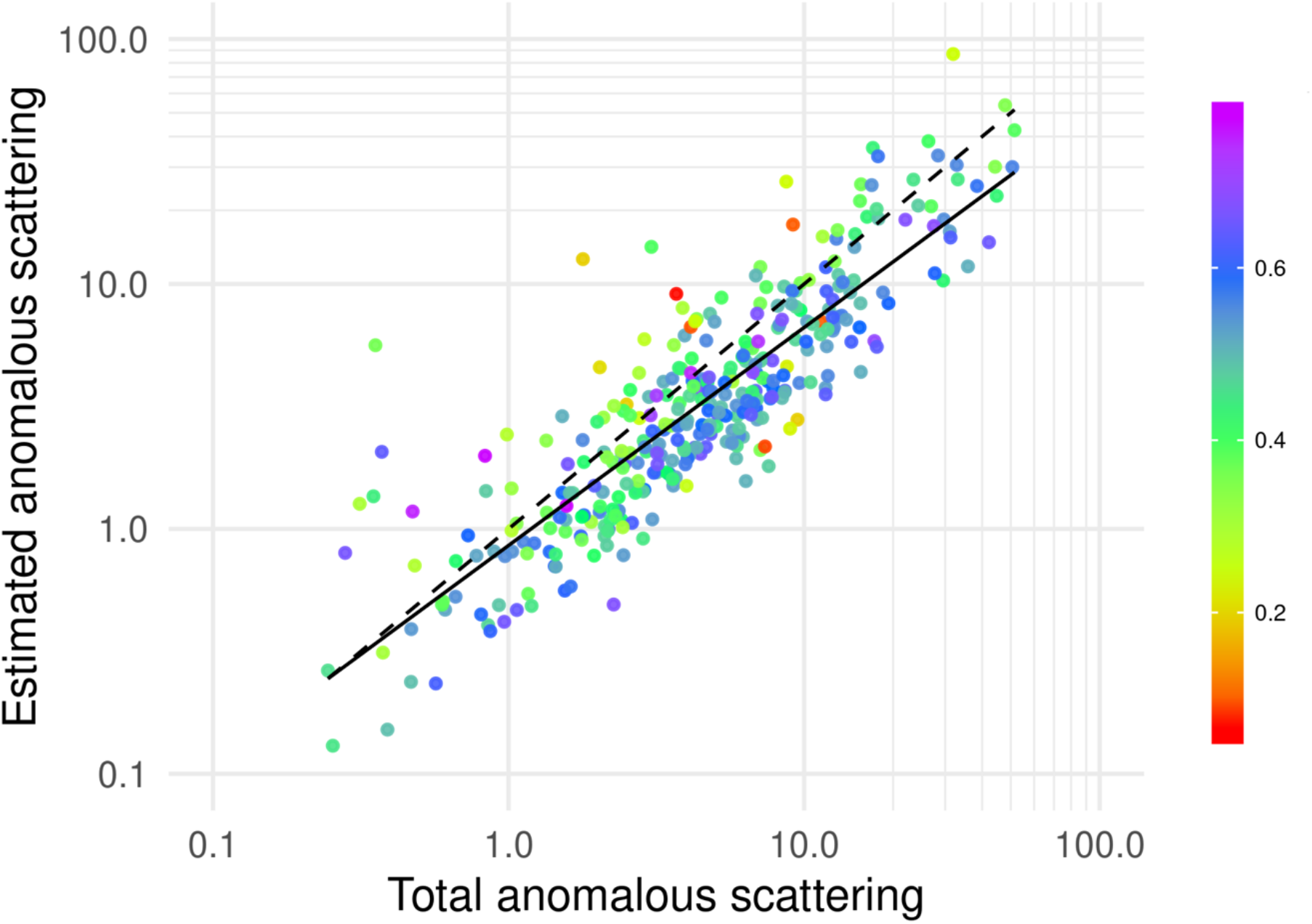
Estimation of equivalent fully-occupied number of primary anomalous scatterers, for data deposited as either intensities or amplitudes. The horizontal axis is the total anomalous scattering power of the gold-standard substructure (f”-weighted sum of squared occupancies of refined sites) and the vertical axis is the estimated anomalous scattering power. The dashed black line represents a perfect prediction while the solid black line shows the least-squares linear fit of the estimates. Each data point is coloured by the map correlation coefficient as shown in the legend. Both axes are plotted on a log_10_ scale.

**Supplementary Figure 2.**
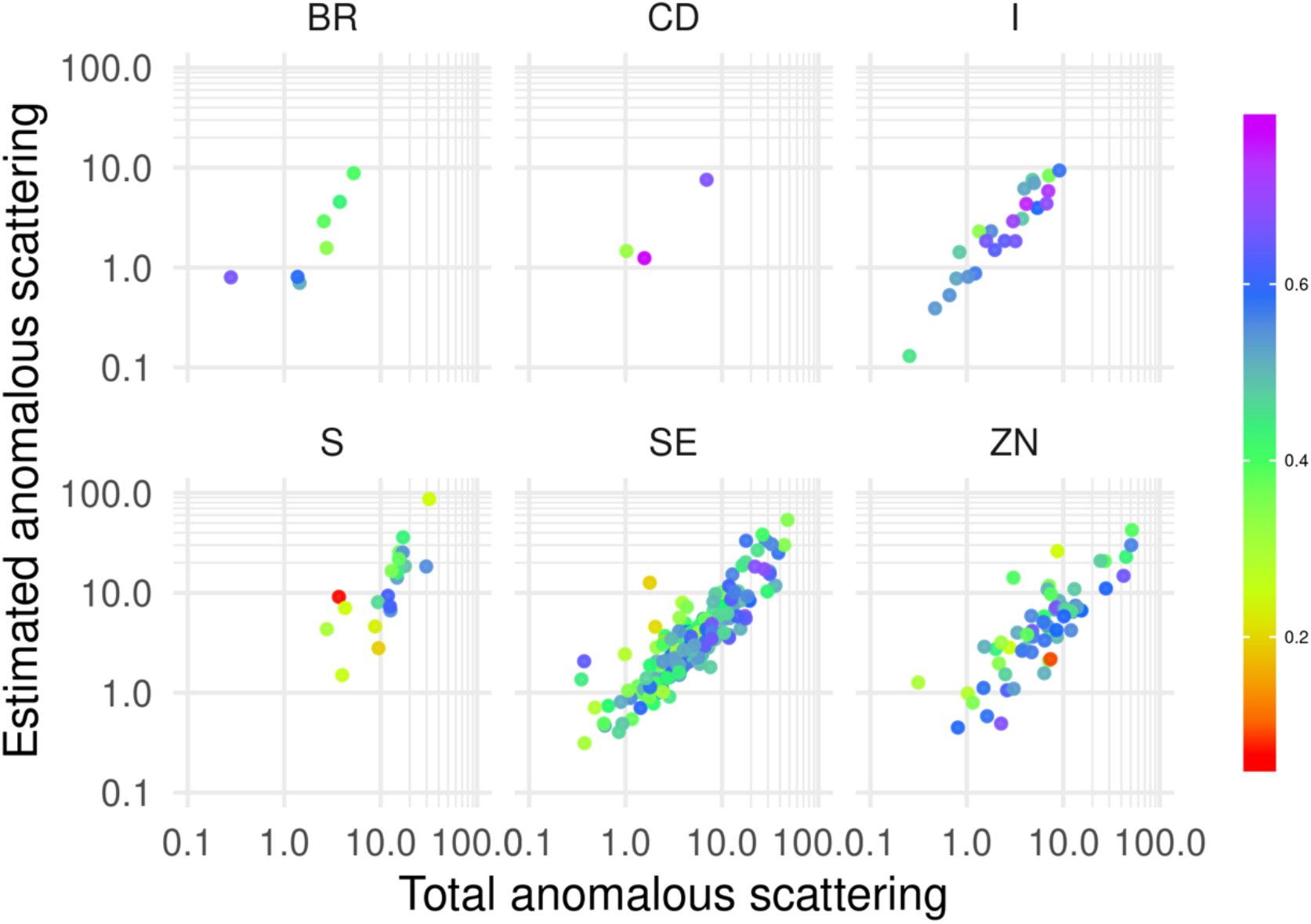
Estimation of equivalent fully-occupied number of anomalous scatterers for different element types. The horizontal axis is the total anomalous scattering (f”-weighted sum of squared occupancies of refined sites) and the vertical axis is the estimated anomalous scattering power. Each data point is coloured by the map correlation coefficient as shown in the legend. Both axes are plotted on a log_10_ scale. Only data for elements with at least 3 cases and over 75% contribution to the total anomalous scattering power are illustrated in this figure.

